# Targeting dendritic cells with RNA-loaded nanoparticles grafted with short peptides

**DOI:** 10.1101/2025.11.05.686747

**Authors:** Vladimir Stamenković, Mislav Brajković, Coral Garcia-Fernandez, Salvador Borrós, Xevi Biarnés, Cristina Fornaguera

## Abstract

Nanoparticles encapsulating therapeutic RNA have emerged as a transformative strategy in precision medicine, capable of mobilizing the immune system to induce specific responses, ranging from immune tolerance to fighting tumors. However, most current preclinical and clinical efforts rely on non-targeted delivery systems, limiting their safety, therapeutic efficacy, and selectivity. To enhance the therapeutic index of RNA-based therapeutic systems for immunomodulatory purposes, we report on the design of a novel Clec9A-targeted polymeric nanoparticle, aimed at selectively engaging dendritic cells responsible for antigen presentation. We began by evaluating *in silico* the binding potential of the previously reported 12-amino-acid WH peptide, known for its high affinity to mouse Clec9A, the human ortholog. Using computational tools, we designed and screened truncated variants of the peptide and identified promising candidates with retained or enhanced binding capacity to human Clec9A. These optimized short peptides were synthesized and covalently conjugated to our proprietary poly(beta amino ester) (pBAE) polymers. We evaluated the impact of conjugation site, comparing terminal versus lateral chain attachment on receptor targeting and confirmed *in vitro* that peptide orientation significantly influences binding efficiency. Additionally, we computationally generated and validated shorter mutant peptide variants with improved Clec9A affinity over the original sequences. Our findings demonstrate that rationally engineered short peptides, when site-specifically conjugated to pBAE polymers, can provide high-affinity, selective targeting of dendritic cells via Clec9A. This strategy lays the groundwork for the next generation of targeted RNA-based immunotherapeutics, offering improved selectivity, immune activation, and therapeutic potential.

**Graphical abstract:** Schematic representation of the workflow used in this work.

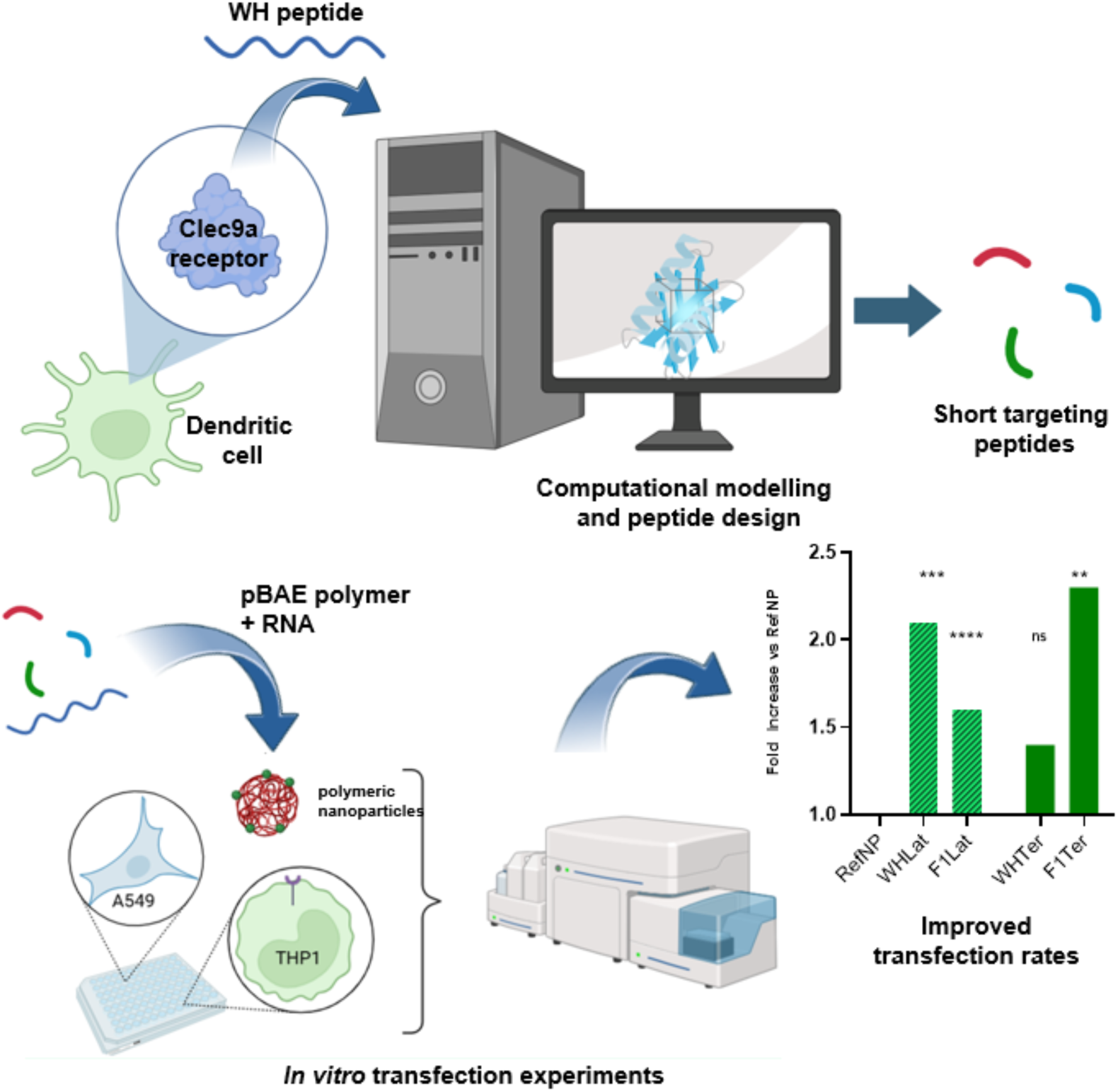

## Introduction

Dendritic cells (DC) are professional antigen-presenting cells (APC) whose primary role is the regulation of T-cell differentiation. In recent decades, many studies have explored the therapeutic potential of immune modulation through DC manipulation^1,2^. A variety of responses can be achieved depending on the type of DC and receptor targeted, and the use of adjuvants^3^. Key molecules for targeting DCs are C-type lectin receptors such as Clec9A, Clec12A, and DEC-205^4^. Clec9A plays a crucial role in extracting and presenting antigens from dead and damaged cells by recognising and binding to actin filaments (F-actin) exposed on them (Figure 1A)^5^. Antigen targeting DC cells via Clec9A can induce strong humoral and cellular responses while avoiding off-target effects^5,6^, even without the presence of adjuvants^3,7^. Possible clinical applications of targeting Clec9A include the induction of transplantation tolerance or controlling autoimmunity^8^, enhancement of vaccine-mediated antibody responses to viral antigens^9,10^, and cancer vaccines^4^.

**Figure 1.**
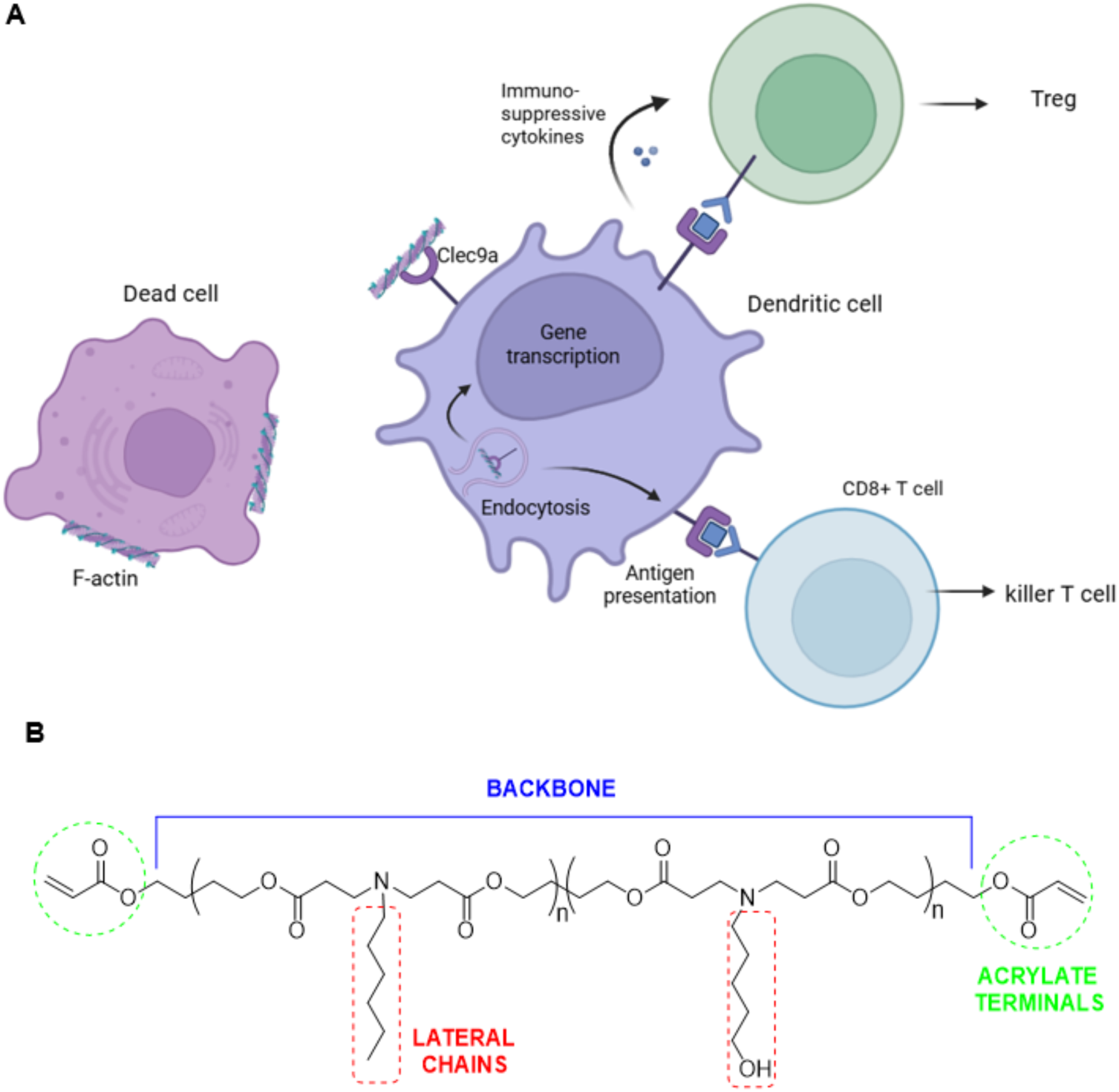
**A)** Recognition of F-actin on dead cells via Clec9a and mechanisms of induction of immune tolerance via differentiation to Treg cells or immune reaction via differentiation to cytotoxic T cells **B)** Complete structure of C6 pBAE polymer and its structural elements

Tumour vaccines have shown enhanced T-cell dynamics and the potential to counteract the immunosuppressive state of the tumour microenvironment, thereby promoting immune cell infiltration. mRNA-based vaccines in particular offer many advantages, such as adaptable design, production scalability, and the possibility of therapy personalization^11,12^. They encode tumour-specific antigens to be translated into proteins in patients’ cells, particularly APC, which then provoke an immune response against the cancer ^11^. Their translation into clinical use, however, is hampered by immune tolerance, weak immunogenicity, non-specific immune system activation, and low efficacy^13,14^. Here, targeting DC presents themselves as an attractive strategy to overcome these problems. A subset of these cells, known as type 1 conventional DCs (cDC1), excel in the cross-presentation of exogenous antigens via MHC class I to naive CD8+ T cells. It has been found that in cancer patients, the abundance of cDC1 in tumour tissue correlates with survival outcomes and the efficacy of immune checkpoint inhibitors^15^.

Besides potential activation of cytotoxic T cells, cDC1s also have immunoregulatory functions and can mediate immune tolerance for overcoming autoimmune reactivity. Use of immunosuppressive drugs can induce tolerance DCs, but their non-specific delivery can reduce overall immunity, thus increasing susceptibility of the patient to infections and other side effects^16^. DCs integrate various immune signals to restore immune homeostasis through a balance of inducing inflammatory T-cell apoptosis, regulatory T cells (Tregs) and modulation of pro- and anti-inflammatory responses. DCs targeted with specific autoantigens have been reported to induce superior tolerogenic responses in comparison with standard immunosuppressive treatments^17^. Combination of autoantigens with tolerogenic agents within a single vehicle can guarantee optimal DC response and stimulation of Treg proliferation^18^. DC targeting strategies have been explored to treat graft rejection^19^, multiple sclerosis ^20^, type 1 diabetes^17^ and systematic lupus erythematosus^21^.

cDC1s are present in both humans (CD141+/BCCA3+) and mice (XCR1+ CD8+ or CD103+), and share functional similarities^4^, important fact in terms of validating novel nanotherapies before going to clinical trials. Among these similarities, both species share equivalent plasmatic membrane surface receptors. These surface proteins are important when aiming at active targeting of the nanosystems to specific cell lineages, such as cDC1. Indeed, recognizing receptors exclusively expressed in cDCs is a must for the design of actively targeted vaccines. Clec9A is expressed consistently and selectively on cDC1 in both species. Mouse Clec9A protein bears 53% structural resemblance to the human Clec9A, including most structural features, such as intracellular signalling motifs^22^. Recently, a short peptide called WH with specificity towards mouse Clec9A was developed^23^. Computer aided docking of the WH peptide to Clec9A identified key contact interactions with Asp248 and Trp250 from the receptor, which were confirmed by mutagenesis. Coupled with OVA257-264 epitope, this peptide successfully activated OVA-specific CD8+ T cells and reduced lung metastases in a B16-OVA lung metastasis mouse model^23^. Nevertheless, molecular interaction between this peptide and the human receptor were not yet studied. Specifically when attached to a nanovehicle, it requires a further study. In this context, the use of mRNA coding specific antigens is advantageous because of their ease of production, adaptability and ability to produce longer lasting immune responses without the need for reformation. In recent years, encapsulation of mRNA has been performed inside of solid lipid nanoparticles, with cationic polymer nanoparticles emerging as alternative candidates^24^.

Among different types of nanosystems, poly(beta-amino ester) (pBAE) based nanoparticles (NPs) present a promising candidate for encapsulation of mRNA^25^. pBAE are biodegradable and biocompatible polymers, capable of forming nanometer-scale structures around genetic material and enhancing its transfection. Oligopeptide end-modified pBAE have demonstrated high efficiency of encapsulating any type of genetic material ^26,27^. Another advantage of these polymers is versatility in design for desired purposes. Modifying pBAE structure (backbone, lateral chains and acrylate terminals) allows for tuneable properties of NPs such as surface charge, hydrophobicity and targeting capacity (Figure 1B) ^28,29^. Until now, it remains unclear how the targeting moiety position on the polymer (attached to lateral chain or acrylate terminals) affects its surface exposure during NP assembly and consequently targeting efficiency.

In this study, we first evaluate the interaction between WH peptide and human Clec9a *in silico* and explore the possibility of designing shorter variants of the peptide to be covalently bonded with the NP building blocks that retain, or even increase, the affinity towards the receptor. Shortening targeting peptides offers improved synthesis yields and reduced costs, ease of conjugation and better receptor accessibility, while reducing immunogenic potential and chemical instability. The designed variants of the WH peptide are then validated *in vitro* by attaching them to different positions of the pBAE backbone and evaluating the NPs’ transfection efficiency on human cells. We show that shorter versions of the WH peptide attached to pBAE acrylate terminals improve transfection equally or better than laterally attached full-length peptide.

## Experimental part

### MATERIALS

Reagents and solvents used for synthesis were purchased from Sigma Aldrich, Iris Biotech and Thermo Fisher Scientific. Peptides used for synthesis of oligopeptide modified poly-β-amino esters (OM-pBAE) – CK3 (CKKK), CH3 (CHHH), WH (WPRFHSSVFHTH), CWH (CWPRFHSSVFHTH), F1 (WPRF), F1C (CWPRF) were obtained from SB peptide (France) with at least 95% purity. F1M (WPAF) and F1MC (CWPAF) were synthesized with Liberty Blue 2.0 Microwave Peptide Synthesizer (CEM Corporation, USA). Addition of 0.1 M HCl was used to replace TFA counterions. mRNAs used in this study were eGFP (CleanCap Enhanced Green Fluorescent Protein mRNA 5-methoxyuridine); obtained from TriLink. Cyanine 5 (Cy5) NHS ester was purchased from Lumiprobe, PE-anti-human-Clec9a antibody was obtained from Biolegend.

### CELL CULTURE

THP-1 (ATCC TIB-202) human monocyte cells were maintained in RPMI-1640 medium supplemented with 2 mM L-glutamine, 1 mM sodium pyruvate, 100 IU/mL penicillin/streptomycin and inactivated fetile bovine serum (FBS) at a 10% final concentration. A549 (ATC CCL-185) human lung adenocarcinoma cells were grown in DMEM medium supplemented with 2 mM L-glutamine, 100 IU/mL penicillin/streptomycin and FBS at a 10% final concentration. All cells were cultured at 37°C, under a 5% CO_2_/ 95% air atmosphere until 90% confluence before starting transfections.

## METHODS

### Modelling Clec9A and WH peptide structures

The human Clec9A structure was retrieved from the Protein Data Bank (PDB ID: 3VPP)^30^. The structure lacked residues 203–208 and 238–241 and included a non-native S225D mutation. To obtain a complete, wild-type model, homology modeling was performed using HHpred ^31^, with human (3VPP) and mouse Clec9a (PDB ID: 3J82)^32^ as templates. The WH peptide sequence was retrieved from ^23^: WPRFHSSVFHTH. Three-dimensional structures were generated using two independent methods: PEP-FOLD ^33^from which the two structures with the lowest energy and highest TM-score were selected; AlphaFold ^34^, from which the first two top-ranked predicted models were used.

### Molecular Dynamics Simulations

All simulations were performed using GROMACS ^35^. Initial protonation states at pH 7.4 were assigned using H++ ^36^. Histidine protonation was defined as follows: HIS135 (δ and ε), HIS182 (ε only); HIS10 in WH peptide was protonated on ε (AlphaFold) or δ (PEP-FOLD), respectively. Disulfide bonds were retained as in the original PDB structure. Simulations used the AMBER99SB-ILDN force field for proteins and TIP3P water model. Each system was solvated in a cubic box with at least 1.0 nm distance between solute and box edge. Systems were neutralized and adjusted to an ionic strength of 150 mM NaCl. Energy minimization was conducted in 6 cycles (5,000 conjugate gradient steps each), with a steepest-descent step every 1,000 steps. Equilibration followed a 3-stage protocol: solvent equilibration: 500 ps (1 fs timestep), 300 K, restrained protein. Heating: 200 ps (2 fs), temperature annealed by gradually increasing from 175 K to 310 to 310 K (Nose–Hoover thermostat ^37^, τ = 2 ps). NPT equilibration: 200 ps (2 fs), 310 K, constant pressure using the Parrinello–Rahman barostat ^38^(τ = 4 ps). Production simulation runs were carried out in multiple replicates in the constant volume and temperature (npt) ensemble at 310 K for 500 ns for both unliganded Clec9a and for each initial Clec9a:WH geometry, and 200 ns for each initial WH peptide structure. MD trajectories were concatenated, and RMSD-based clustering was performed with GROMACS tools following ^39^. A RMSD cutoff of 0.21 nm was used for Clec9a systems and 0.45 nm for the WH peptide alone. Representative conformations of Clec9a (most populated cluster) and WH peptide (top two clusters from initial PEP-FOLD and AlphaFold geometries) were selected for docking.

### Molecular Docking (MD) Clec9A:WH

Docking simulations were performed using AutoDock4 ^40^. The Clec9 receptor was kept rigid; the WH peptide ligand was fully flexible. Blind docking was performed by using multiple grid boxes (n = 8) defining different search spaces spread along the Clec9 structure. The genetic algorithm was followed with a population size of 300 for 300 runs using exhaustive search parameters. Ten representative Clec9A:WH structures of the complex were selected for MD based on: high population within the docking results (clustering with RMSD cutoff = 3 Å) and low docking energy within the cluster.

### Contact Analysis

Four Clec9A binding sites were defined based on visual inspection of MD trajectories: Site 1: residues 114–231, Site 2: residues 120–220, Site 3: residues 178–226, Site 4: residues 132–227. Time-resolved contact analysis was performed for each site using GROMACS tools. Normalized contact profiles were computed by dividing contact counts by the site-specific maximum. From normalized data, residence time analysis was derived by identifying the dominant binding site at each frame. Relative occupancy was calculated across all frames and replicates

### WH Peptide Fragments and Mutants

Three WH peptide fragments were designed: WH14 (residues 1–4: WPRF), WH58 (residues 5–8: HSSV), WH912 (residues 9–12: FHTH). Each fragment was modeled in four separate Clec9a-bound systems (one per binding site). From representative bound conformations, extraneous residues were removed to generate minimal complexes. WH14 mutants were generated via single alanine substitutions (APRF, WARF, WPAF, WPRA) by modifying side chains in the PDB files. These systems were subjected to MD and contact analysis as described above for the full-length peptide.

### Synthesis of OM-pBAEs

The backbone of all oligopeptide-modified poly-β-amino esters was the so-called C6 pBAE. Synthesis of the backbone and subsequent attachment of cysteine-containing oligopeptides (CK3, CH3, CWH, CF1, CF1M) to acrylate terminals via thiol-Michael addition were performed as previously described ^26^. Briefly, C6 polymer was synthesized by measuring hexylamine, 5-amino-1-pentanol, and 1,4-butanediol diacrylate at amines/diacrylate 1:1.2 molar ratio in a round-bottom flask and heating the reaction mixture at 90°C overnight under constant stirring. The obtained product was then cooled down to room temperature and kept at -20°C until further use, without purification. For thiol-Michael addition, a corresponding amount of peptide and C6 pBAE at a 2.1:1 ratio was prepared in anhydrous DMSO and stirred at 25°C overnight. The product was purified by precipitation over a cold diethyl ether/acetone mixture at 7/3 (v:v) ratio and two subsequent washes with the same solvent mixture. Lateral chain labelled OM-pBAEs were prepared by Steglich esterification of the hydroxy group in the lateral chain of the C6 polymer. C6 pBAE and the peptide (1.1 eq) were dissolved in anhydrous dichloromethane/dimethylformamide 9:1 (v:v) mixture, after which dicyclohexylcarbodiimide (DCC, 1.1 eq) and 4-dimethylaminopyridine (DMAP, 0.1 eq) were added, and the reaction proceeded under constant stirring at 25°C overnight. At the end of the reaction, the mixture was passed through through a 0.45 μm nylon filter to remove the precipitate, most of the solvent was removed under vacuum, and then the product was precipitated and washed with ether/acetone mixture 7/3 (v:v) mixture and dried under vacuum. Subsequently, the product’s acrylate groups were end-capped with CH3 as described above.

CH3 end-capped C6 pBAE was labelled with Cy5 NHS ester by preparing a 10 mg/mL stock solution of Cy5 NHS in DMSO, and adding it to a DMSO solution of CH3 C6 pBAE (10 mg/mL) with triethylamine at 1:1 molar ratio and incubating the mixture for 20h at room temperature. The resulting labelled polymer was precipitated and washed with 7/3 (v:v) diethylether/acetone mixture. The product was redissolved in DMSO at 100 mg/mL concentration and kept at -20°C until use. Labelled H polymer was diluted with unlabelled polymer to obtain a mixture for preparing NPs for uptake experiments, adjusting the concentration by measuring fluorescence intensity of the mixture dilution in phosphate buffered saline (PBS). Synthesized structures were confirmed by 1H-NMR. NMR spectra were recorded on a 400 MHz JEOL (JEOL Ltd., Japan). All data was processed and analysed using the MestreNova 12.0.3 software. Solvents used were CDCl_3_, and DMSO-d6.

### NPs preparation

Briefly, each NP formulation was prepared at a OM-pBAE: nucleic acid ratio of 25:1 w/w, by mixing equal volumes of GFP mRNA (eGFP, concentration 0.5 mg/mL) with a corresponding mixture of pBAE polymers in 12.5 mM AcONa buffer solution. RefNP, as control non-targeting formulation contained 60 % of CK3 OM-pBAE and 40% of CH3 OM-pBAE, while the targeting formulations contained 60% of CK3 pBAE, 30% of CH3 pBAE and 10% of corresponding targeting peptide OM-pBAE. In the case of uptake experiments, CH3 OM-pBAE labelled with Cy5 was used instead of regular CH3 OM-pBAE. eGFP was added over the polymer solution and mixed by pipetting, followed by incubation for 30 min at 25°C. Afterwards, the complexes were nanoprecipited in an equal volume (Vi) of DEPC water, and diluted in another equal volume (Vi) of HEPES 20 mM + 4% sucrose. The NPs were then lyophilized and kept at -20°C until further use.

### Physicochemical characterization of NPs

#### Dinamyc light scattering (DLS)

Hydrodynamic diameter (z-ave), polydispersity index (PdI) and zeta potential (ζ) were measured at 25°C, using a ZetaSizer Nano ZS (Malvern Instruments Ltd, United Kingdom), with a 633 nm laser at a 173° detection angle. For size measurements, NPs of the initial nucleic acid concentration were used, while 1/100 dilution in deionized water was prepared for measurements of zeta potential. Three measurements of each batch were performed with 20 runs per measurement, taking into account intensity approximation. The particle size of the complexes were additionally determined by NP Tracking Analyzer (NS300 NTA equipment, Malvern Instruments Ltd, United Kingdom), using a 488 nm laser. Measurements were carried out at 1/100 dilution in DI water. Three measurements of at least 30 s and a total number of 80-100 NPs per frame were analysed. Results were plotted as mean and standard deviation of triplicate analyses by intensity.

#### Encapsulation efficiency (EE)

Encapsulation efficiency was evaluated by electrophoretic mobility shift assay. Each batch was added over agarose gel (2.5% agarose w/v) in Tris-Acetate-EDTA (TAE) buffer containing ethidium bromide (1 µg/mL). Finally, 10 µL of sample was added to the wells and the electrophoresis was performed for 1 hour at 80V (Apelex PS 305, France) and mRNA bands were visualized by UV radiation. Encapsulation efficiency was quantified using a Ribogreen kit. NPs were freshly prepared and mixed with Tris-EDTA (TE) buffer and Ribogreen dye. Three standard curves were prepared. First standard curve was a mRNA calibration curve ranging from 1.125 to 22.5 μg/mL in TE buffer. Second standard curve was a nucleic acid: NPs calibration with heparin in the same range of concentrations. Third standard curve was a heparin calibration. Samples were evaluated both in the presence and absence of heparin and encapsulation efficiency was calculated as follows:

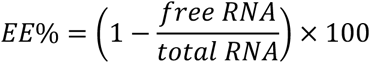

### Confocal microscopy

For imaging of THP1 cells, 3 × 10^5^ cells per well were seeded into 24-well plate. Cells were then fixed with a 4% (w/v) paraformaldehyde (PFA), permeabilized with 0.3 % Triton X-100 in PBS and blocked in 2% FBS in PBS. Nuclei were stained with DAPI, while PE mouse-anti-human Clec9A was used for immunostaining. Approximately 50 µL were added to the surface of coveslips and Fluoromount-GTM mounting medium was added. Samples were finally covered with slides and edges of cover glass sealed with nail polish. Images were captured with a confocal laser-scanning fluorescence (CLFM) Leica SP5 (Molecular Probes, Leica Microsystems, Germany). Images were processed in ImageJ software.

### Transfection experiments

Cells were seeded in a 96-well plate at 10000 cells/well to a roughly 90% confluence 24 hours prior to starting the experiment. Cells were incubated at 37°C, under a 5% CO_2_/ 95% air atmosphere with 0.6 µg/mL of eGFP mRNA encapsulated within pBAE polyplexes. Experiments were stopped after 24h of incubation, fixed with 1% PFA and analysed for GFP expression by flow cytometry (Novocyte, ACEA Biosciences Inc., USA). GFP expression was compared against non-treated cells as negative controls and Lipofectamine2000 as a positive control. There were three wells for each condition, and each experiment was repeated in triplicate. For competitive transfection assay, two 96-well plates with the same conditions were prepared – one was incubated in a fridge at 4°C and the other in an incubator at 37°C for 1h. Afterwards, NPs were removed from both plates and fresh medium introduced in each well and the experiment continued until 24h since its start. For cellular uptake experiments, cells were incubated with pBAE-eGFP polyplexes formulated with Cy5 labelled H polymer at 0, 18 and 22h timepoints.

### Statistical analysis

GraphPad Prism was used for statistical analysis. Triplicate of experiment results were joined into one graph and analysed for each transfection experiment. One-way ANOVA was used to determine whether there was difference across formulations. If statistical significance was found, Dunnett’s post hoc test was performed to compare each formulation against RefNP as control. Two-way ANOVA was used to determine significant differences in competitive assay. If statistical significance was found, t-test was performed to determine differences between temperatures. *p* value < 0.05 was considered statistically significant. Abbreviations used in the manuscript: ns = no significant differences; \**p* < 0.05; \*\**p* < 0.01; \*\*\**p* < 0.001; \*\*\*\**p* < 0.0001. Figures were designed with Biorender software.

## RESULTS AND DISCUSSION

### Modelling human Clec9A – WH peptide complex

The WH peptide was originally designed *in silico* and demonstrated specific binding towards the mouse Clec9A receptor^23^. However, the binding of the same peptide to human Clec9A is to be assessed. Since binding of this peptide onto Clec9A surface is mediated by weak protein-protein interactions, special attention is to be given to protein flexibility and explicit solvent effects. For this reason, we modelled the three-dimensional structure of human Clec9A in complex with the WH peptide by means of artificial intelligence (AI) model generation, MD and evaluated its stability by molecular dynamics simulations. The structure of the extracellular domain of human Clec9A receptor was taken from experimental crystallographic data and missing elements modelled by homology modelling. The resulting Clec9A structure was equilibrated in solution by long molecular dynamics simulations. The most representative structure of Clec9A was taken to predict the binding of WH peptide (see Figure 2A). Given the short size of the WH peptide (12 amino acids), it is not expected to adopt a single defined three-dimensional structure in solution. Starting structures of the WH peptide were generated with ALPHAFOLD and PEPFOLD. These rendered a random coil conformer and alpha-helix geometry respectively. An ensemble of conformations of the WH peptide were generated in solution by running two replicate molecular dynamics simulations starting from each initial structure. Trajectories were clustered and the four most representative conformations of the WH peptide were selected for binding to Clec9A. The selected structures adopt alpha-helix, beta-sheets and random coil conformations (see Figure S1). The three-dimensional structure of the complex between each peptide conformer and the modelled human Clec9A structure was predicted by MD. Different docking set-ups were used to ensure a proper exploration of the whole Clec9A protein surface. The whole ensemble of putative Clec9A-WH structures was collected from the dockings. These were ranked by the binding affinity score provided by AutoDock ^41^. Complexes were clustered by structure similarity and 12 representative high-affinity Clec9a-WH structures were selected for further validation (Figure 2A). The solvent and thermal stability of WH peptide on the surface of Clec9A at these initial positions was further assessed by molecular dynamics simulations. Twelve different simulation replicas were performed, and all simulation data was gathered and analysed together. Several unbinding and rebinding events of the peptide were observed along the trajectories, reflecting a distribution between bound and unbound states at different sites of the protein structure. The simulations revealed that there are 4 putative binding sites along the Clec9A surface on which the WH peptide is stable enough. The residence time of the WH peptide bound at each subsite was monitored in all replicas (Figure 2B). Overall, the WH peptide mainly resides in binding sites 1 (29% of the time) and 2 (31% of the time), whereas residence in binding sites 2 and 3 is lower. The peptide is not bound to any of these sites 6% of the time, which reveals that, despite not being site-specific, the WH peptide strongly adheres to the Clec9A surface. Overall, given the sequence and structural similarity of human and mouse Clec9A, and the stability of the complex exhibited by molecular dynamics simulations, we assume that human Clec9A can also be targeted by the WH peptide.

**Figure 2.**
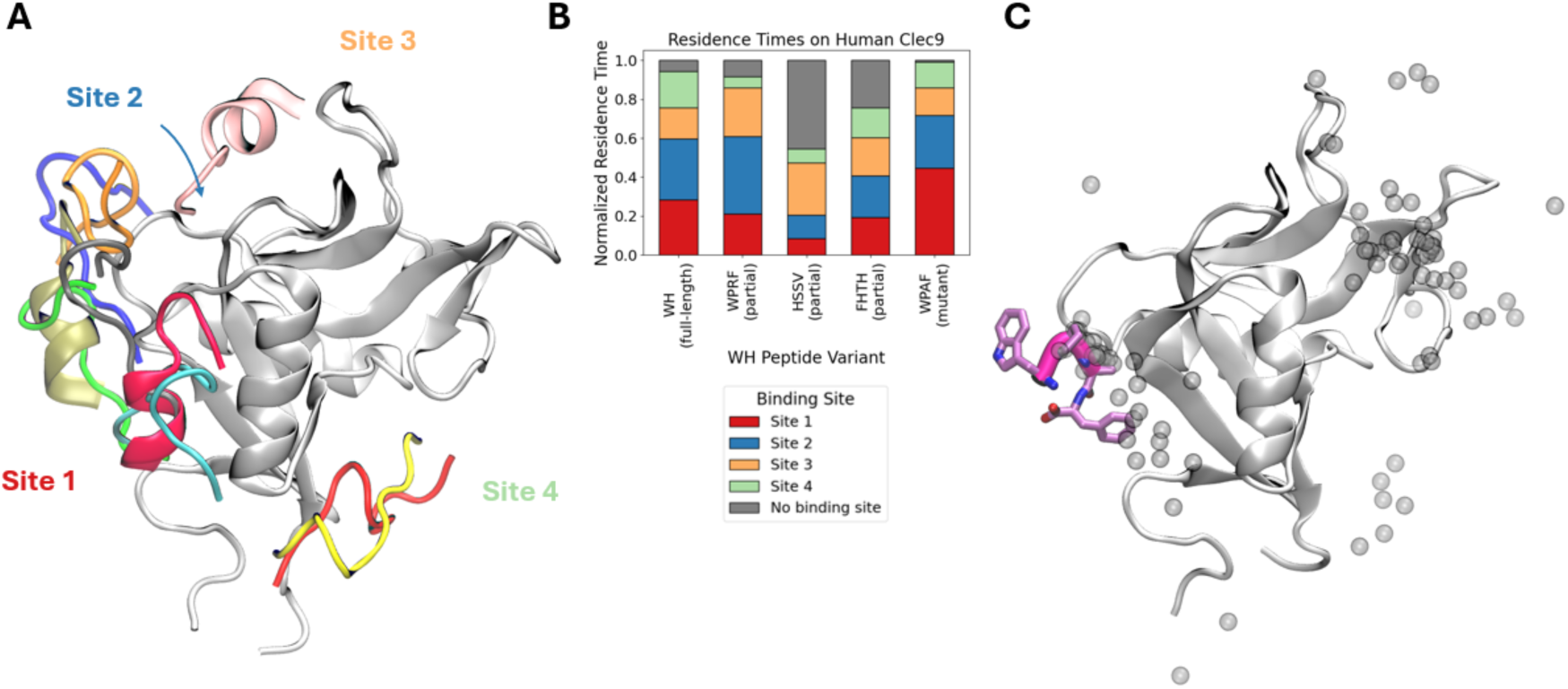
Modelled structures of human Clec9 in complex with WH peptide and variants. Protein backbone coloured according to putative WH binding sites. **A)** Superposition of representative Clec9A:WH geometries obtained via computational docking. **B)** Residence time of WH peptide and its variants at each binding site. **C)** Preferential binding mode of WPAF on Clec9 structure at Site 1 obtained by molecular dynamics (shown in magenta). Alternative binding modes explored during the dynamics are shown in grey spheres.

### Design of truncated forms of the WH peptide

Despite being a short peptide, WH may interfere with the synthesis of the pBAE polymers and the further assembly of the NPs that may compromise the efficient encapsulation of the genetic material. To note that any modification performed on the pBAE polymers will alter the distribution of the cationic charges. Specially, when intending to modify the end terminus of the polymers (i.e. the cationic polymers substitution by these peptides), the cationic charge of the polymer for the RNA encapsulation may be strongly compromised. For this reason, it was envisioned to design even shorter variants of the WH peptide.

Initially, a spare distribution of truncated forms along the WH sequence was considered: WPRF, HSSV and FHTH. The stability of these short variants was assessed again by molecular dynamics simulations. The starting structures were taken from the equilibrated form of the full length WH peptide at each of the 4 early identified binding sites. The three truncations were performed at each of the 4 binding sites, thus leading to a total of 12 initial structures for the different WH truncated peptides in complex with Clec9A. Simulations were performed under the same conditions as for the full-length peptide. In this case, 4 simulation replicas were run for each truncated peptide. The residence time of each peptide at each binding site was measured. Simulations reveal that the WPRF peptide enhances binding at binding site 2 (40% of the time), while binding to other subsites is decreased as compared to the full length WH peptide (see Figure 2B). The HSSV variant displaces binding to binding site 2, although with low frequency (27% of the time) as it remains unbound from Clec9A 45% of the time. Finaly, FHTH variant does not show preferential binding to any site, and it is unbound from Clec9A 25% of the time. Thus, the only truncated variant of the WH peptide that maintains effective binding to Clec9A is WPRF, suggesting that it still could be used as a targeting vehicle.

### Synthesis of the polymers, preparation and characterization of NPs

The proposed polymers that were synthesized are summarized in Figure 3A and a synthesis scheme can be found in Supplementary Information (Figure S2). To laterally attach each of the peptides, the Steglich reaction was performed to form an ester bond between the carboxy group of the C-terminal and the hydroxy group of the lateral chain. NMR spectra confirmed the appearance of peptide signals in the pBAE backbone (Figure S3). In this case, after successful reaction, the acrylate terminals were attached with a CH3 peptide. The same type of thiol-Michael reaction was used to attach cysteine-terminated targeting peptides to the polymer as well. In the NMR spectra, reaction success was confirmed by the disappearance of the acrylate signals between 5.8 and 6.4 ppm. Once the polymers were obtained, formulation of NPs was approached. ^19,34,35^

**Figure 3.**
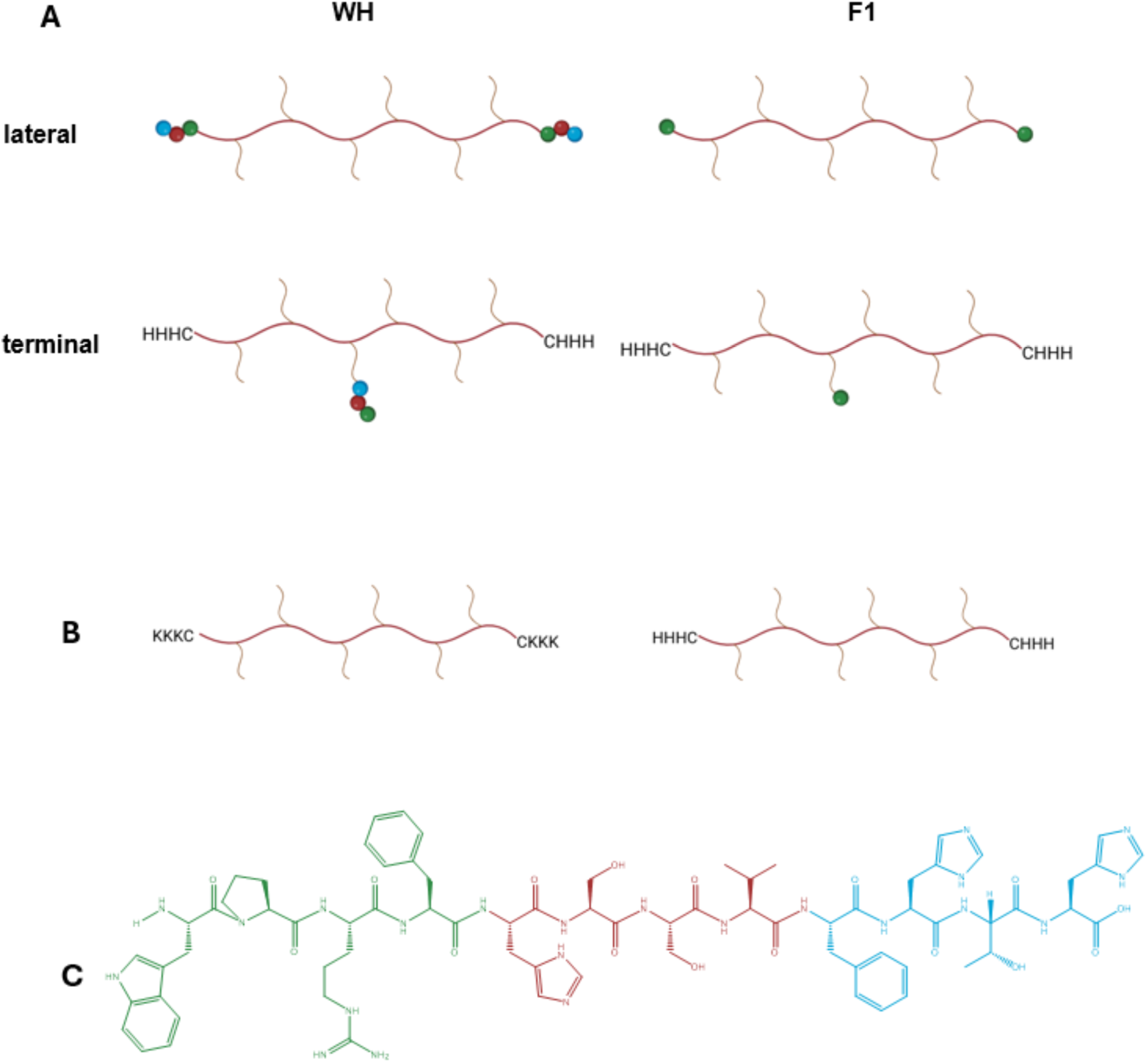
**A)** Graphical representation of the synthesized targeting pBAE polymers with attached peptides **B)** Graphical representation of constituent polymers of the NPs – OM-C6-CK3 on the left and OM-C6-CH3 on the right **C)** Structure of the WH peptide, marked with different colours according to the fragment (green - Fragment 1 (F1) – WPRF, red - Fragment 2 – HSSV, blue – Fragment 3 – FHTH).

The prepared formulations were then submitted to physicochemical characterization in terms of size, polydispersity index, zeta potential and encapsulation efficiency. As it can be seen in Figure 4A, all polyplexes formed have a size of around 180-220 nm, with the reference formulation having an average of 205 nm. When comparing the sizes, a statistically significant difference between RefNP and terminal targeting formulations was determined. While lateral targeting formulations maintain CH3 in their terminals, terminal targeting particles replace a percentage of CH3 with a different peptide, which can explain this difference. WH terminal peptide, as the longest peptide, gives the largest particles in size of the group. Due to its length, it is possible that it influences interaction with genetic material and folding of the peptides, and consequently particles assemble in a slightly different way. All formulations have a low polydispersity index (PdI <0.2), which allows to define them as monodisperse. Significant differences are observed in the same two formulations for zeta potential results. Referent formulation, lateral peptide formulations and F1Ter have similar values of around 36 mV, while WHTer presents lower values. Even though WH contains three histidine residues, the overall effect of the peptide is slight reduction of zeta potential.

**Figure 4.**
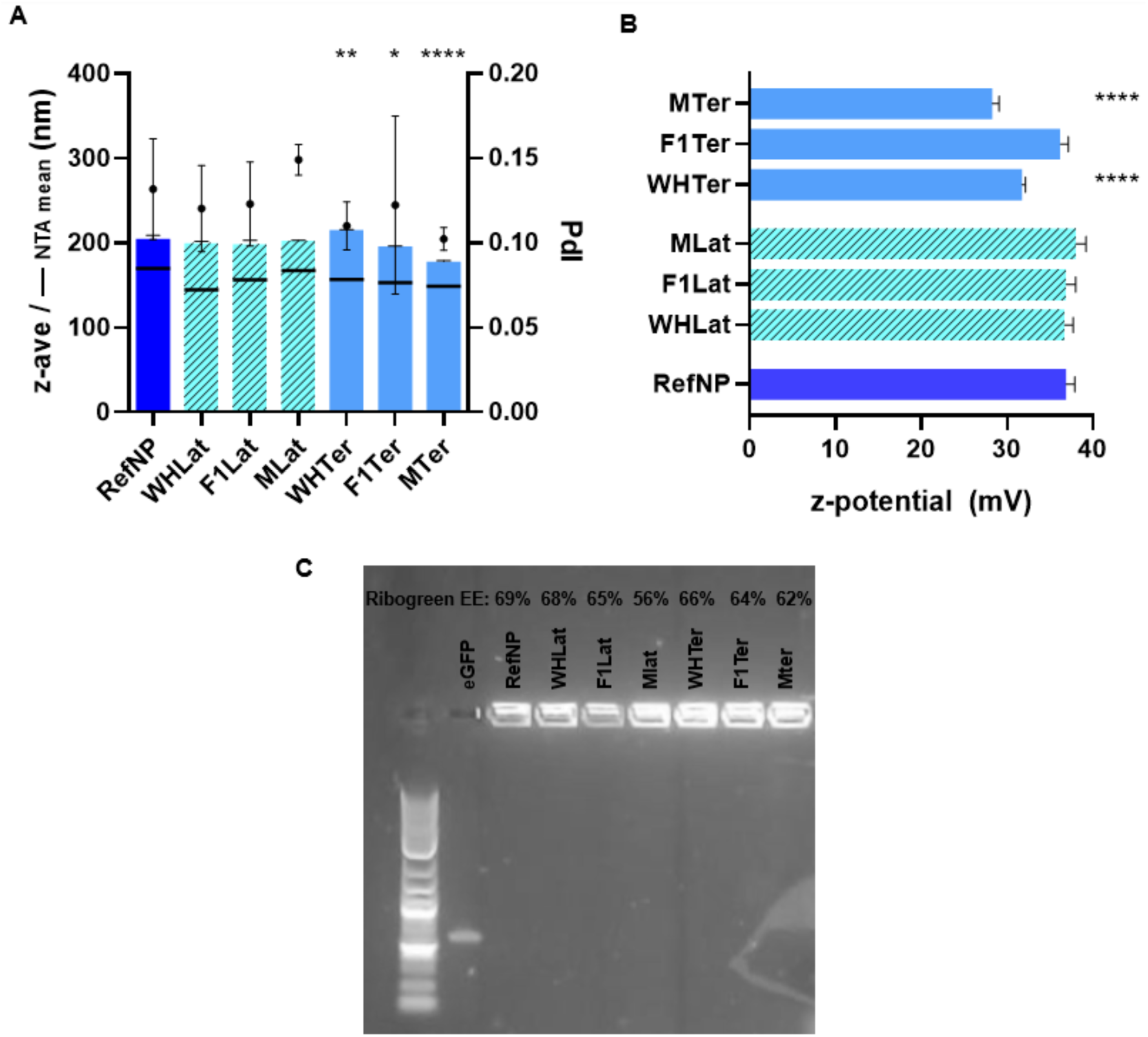
**A)** Results of particle size analysis using DLS and NTA techniques. Hydrodinamic size (z-ave) is presented with a bar graph, while NTA mean size is presented by a line. On the right side of the y-axis, polydispersity index values (PdI) are displayed, each result represented with a dot. **B)** Zeta potential values of the formulations **C)** Agarose gel image after electrophoresis and EE values above each formulation obtained by Ribogreen assay

In the agarose gel retardation assay, all formulations have genetic material that is retained at the starting line, without any additional bands running inside the gel (Figure 4C), which indicates a high degree of encapsulation of eGFP. This method however, is only qualitative and only gives an initial idea of the encapsulation efficiency. To validate these results, the Ribogreen assay was performed to quantify the percentage of the genetic material encapsulated in the NPs (Figure 4C). EE of all formulations was confirmed to be high (between 60 and 70%) in all the prepared formulations. This indicates that in most cases, targeting peptides do not significantly affect the complexation of mRNA.

### *In vitro* transfection experiments

Once all the formulations were validated in terms of size and polydispersity, and encapsulation efficiency, their transfection efficiency was next evaluated. Two cell lines were used. Human epithelial carcinoma cells, A549 cells, were selected as a highly permeable and permissive cell line for transfection studies, without the targeted receptor. THP1 cells, human monocytes isolated from an acute monocytic leukaemia patient, were selected as a restrictive cell line expressing Clec9A receptor. The presence of Clec9A on the surface of THP1 cells was confirmed by confocal imaging when incubated with PE-labelled anti-human Clec9A antibody (Figure S4). These cells are extremely plastic and are easily differentiated into macrophages or mature dendritic cells ^42,43^. Firstly, to confirm interaction between human Clec9A and WH peptide and whether F1 maintains receptor affinity, these two peptides were evaluated on both polymer positions.

Transfection results are presented in Figure 5, and as expected, all NP formulations show a high degree of unspecific transfection for A549 cells with no statistical difference between them. In case of RefNP, as well as targeted NPs, the majority (80-90%) of cells expressed GFP at the end of the experiment, indicating no participation of neither of the peptides in the transportation of the particles through the membrane. On the other hand, THP1 cells are more difficult to transfect and exhibit lower transfection degrees, which is already documented in literature ^44^. Interestingly, our NPs showed differences in transfection across formulations. It can be seen that all four targeting formulations with WH peptide and variants transfect better than RefNP, two of them over 2-fold (WHLat and F1Ter). First of all, this initially confirms computational results of WH-human Clec9A interaction – WH peptide indeed interacts with human Clec9A. Secondly, it was confirmed that F1 – a shortened version of WH does indeed maintain its affinity towards the receptor. To the best of our knowledge, this is the first time in which selective targeting is achieved with such a rationally designed short peptide in NP formulations. Although such a short peptide is not expected to strongly bind the receptor (indeed, dynamic binding is observed during the MD simulations), the local high concentration of peptides on the surface of the NP presumably increases the overall affinity to the receptor via multimerization of the receptor of avidity.

**Figure 5.**
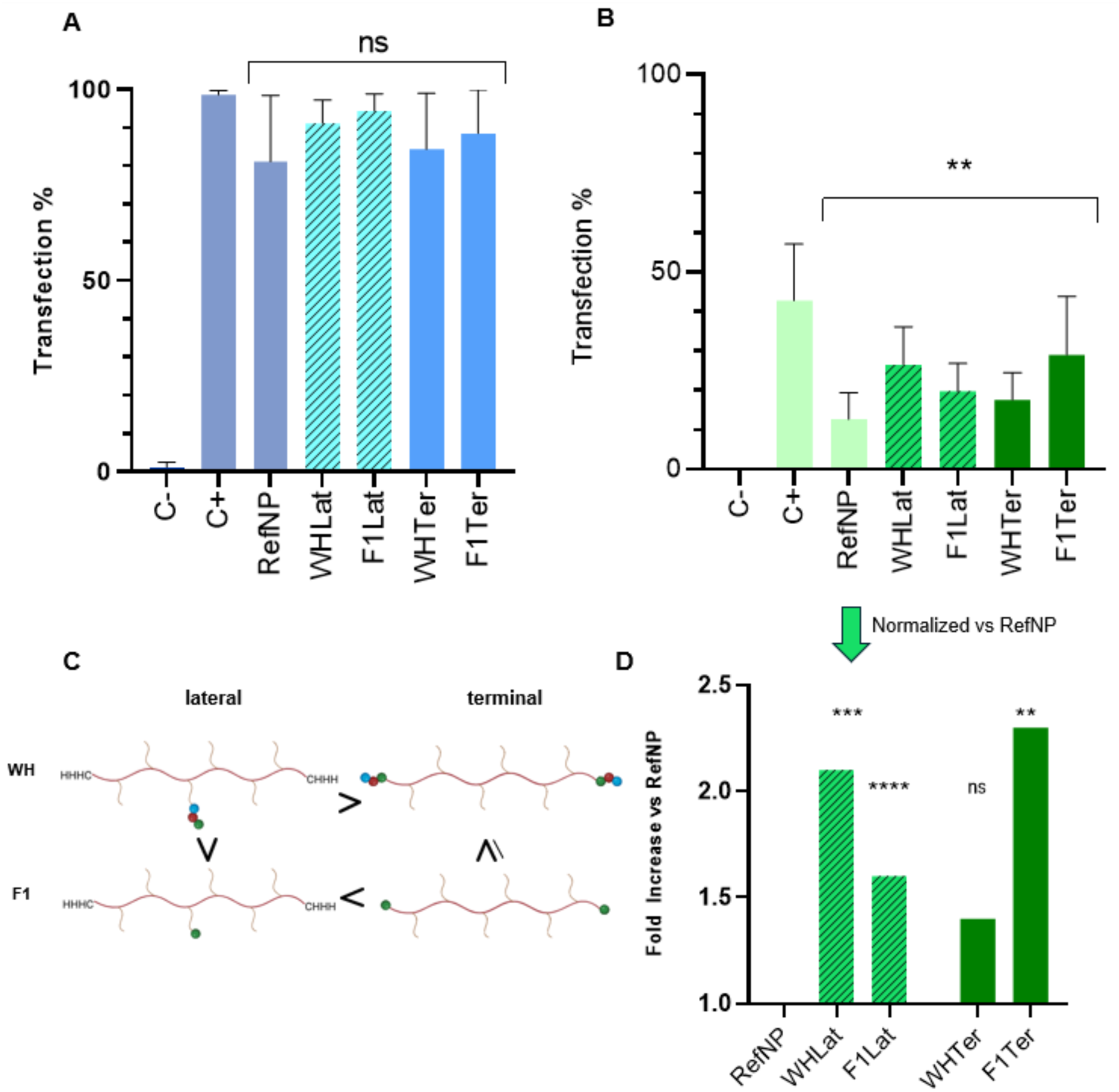
**A)** Transfection results on **A)** A549 (left, blue) and **B)** THP1 (right, green) cell lines; C-negative control, C+ lipofectamine control; **C)** Schematic representation of effect of targeting peptides WH and F1 positioning in C6 pBAE polymer on transfection rates below; **D)** Transfection rates normalized and statistically compared against control formulation RefNP, presented as fold increase

When performing a t-test to compare RefNP against each of the formulations, three of them showed very high statistical significance, with only WHT not being significant. Comparing across polymer positions, it can be observed that WH peptide in the lateral position transfects better than when positioned terminally but also transfects better than laterally attached F1. Interestingly, in the case of F1, the situation is reversed – when positioned terminally, it transfects better than laterally placed, and also than the terminal WH peptide. Terminal F1 transfects equally or better than lateral WH. All these comparisons are schematically summarized in Figure 5C. In conclusion, in the case of pBAE NPs, predicted peptide receptor affinity alone is not a determining factor in the targeting capacity of the final polyplexes, but is also governed by the position of attachment of the peptide to the pBAE polymer. These results highlight the need of having a detailed knowledge of the targeting moiety properties before approaching polymer design. Several properties can be taken into consideration, including hydrophobicity, and in the case of peptides, charged residues and total charge at different pH values. For the peptides used in this study, these characteristics are presented in Table 2.

**Table 2.**
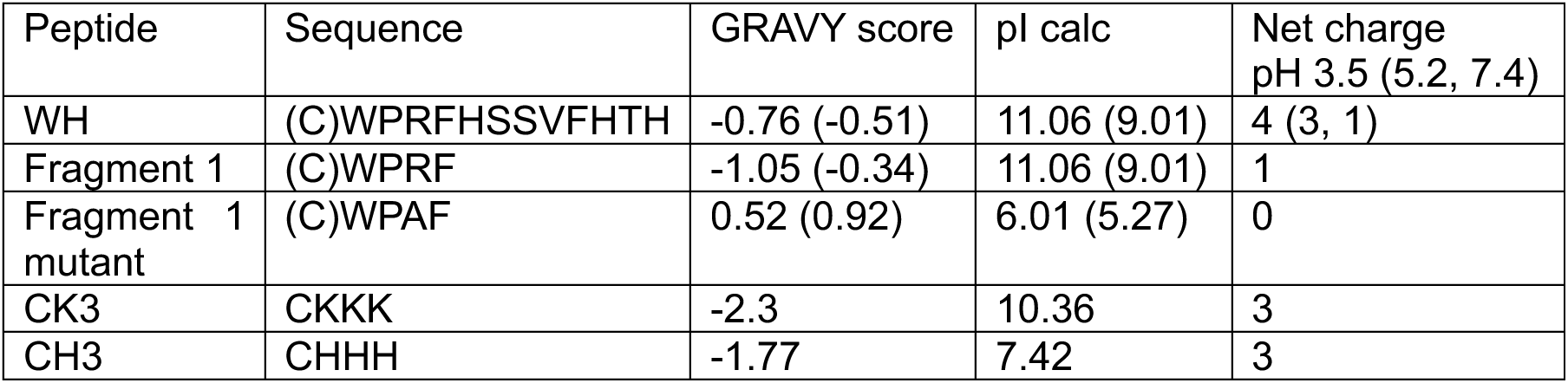
Properties of peptides attached to C6 pBAE polymer, used in this study. GRAVY score was calculated based on the Kyte-Doolittle scale ^45^ . The values in parentheses for GRAVY score and pI values refer to the same peptide containing cysteine (used to attach to acrylate terminals).

In this context, it is interesting to consider the case of the terminally attached WH peptide, which is the targeting formulation that has the lowest success of transfection, which was additionally not found to be statistically significant compared to RefNP. The WH peptide contains three histidine residues that are protonated at particle formation pH (5.2) and could electrostatically interact with the genetic material’s phosphate groups, leading to higher internalisation of the peptide. In a previous study of oligopeptide (CH3, CK3, CR3 and CD3) modified pBAE from our group, it was determined by Förster Resonance Energy Transfer (FRET) that CH3 is relatively closer/more internalised than other oligopeptides ^46^. Additionally, histidine and other aromatic amino acid residues (WH contains two phenylalanines and a tryptophan) can further contribute to interaction with nucleic acids via π-π interactions ^47–49^. Another property to consider is peptide hydrophobicity, which can be expressed with a GRAVY score. Hydrophobicity in the context of NP formation can determine the interaction between hydrophilic genetic material, but also with the more hydrophobic cell membrane.

To confirm that improved transfection percentages of targeting formulations are a consequence of active transport of NPs through the membrane due to ligand-receptor interactions, a receptor-mediated transfection assay was performed. For this, transfection was compared after 1h of uptake of NPs at two different temperatures – 4 and 37°C. At 4 degrees, endocytic uptake pathways are not favoured due to a change in phase state of the cell membrane ^50^ and only passive transport of particles is taken into consideration. Clec9a itself plays a crucial role in antigen processing and presentation via endocytosis ^22^. Therefore, a significant difference in transfection degree in THP1 cells between the two temperatures for a formulation indicates considerable participation of the active mode of transport of NPs, presumably via Clec9A.

Receptor-mediated transfection assay results for A549 cells, similarly to a normal transfection experiment, show no significant difference between formulations at the chosen temperature conditions (Figure 6). In the case of THP1 cells, two-way ANOVA indicates a high statistical significance of the effect of both formulation and temperature, formulation being the most influential factor (70% of total variation). Analysing individual formulations, all targeting formulations show around a 2-fold increase in transfection rates at 37°C in comparison with 4°C, three of them being statistically significant. Thus, active targeting of Clec9A by the NPs decorated with WH peptides was confirmed. The only formulation that is not statistically significant is WHT, which was to be expected given that it gives the lowest degree of improvement of transfection.

**Figure 6.**
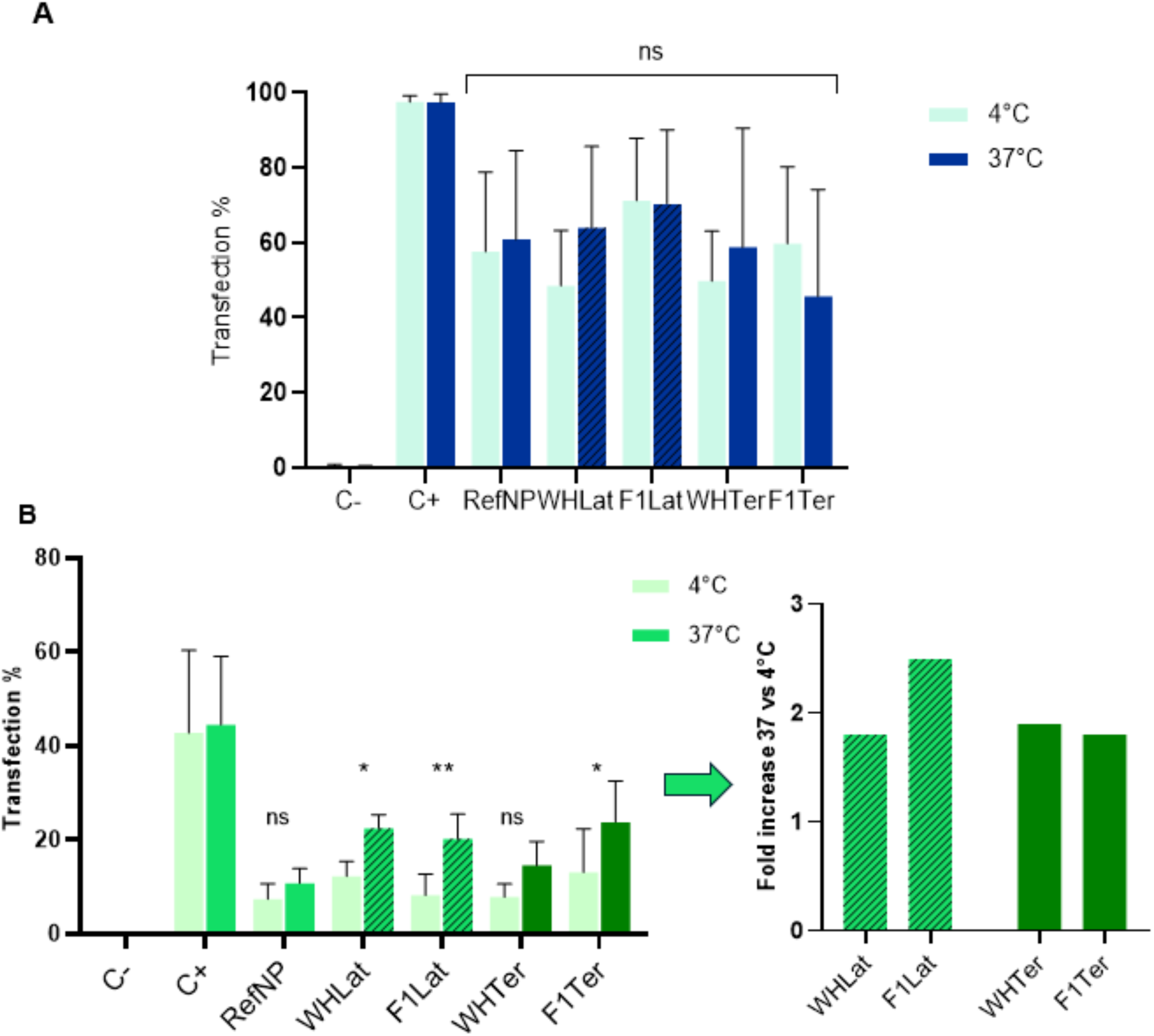
Receptor-mediated uptake assay results at different temperatures for **A)** A549 cell line and **B)** THP1 cells; these results are also presented as fold increase of transfection rate at 37°C against 4°C for each targeted formulation on the right graph

### Modelling of a truncated WH mutant to improve affinity towards Clec9A

Prompted by this successful result, we wondered if binding could be further improved by introducing point mutations to alanine along the sequence. The new variants considered are APRF, WARF, WPAF, WPRA. The stability of these variants on the surface of Clec9A was simulated by molecular dynamics following the same approach. In this case, 16 replicas were simulated in total (4 binding sites for each of the four mutants). Residence time analysis revealed that APRF, WARF and WPRA retained the anchoring to binding site 2 as the original WPRF peptide (see Figure S5). Surprisingly, the WPAF mutant yielded a peptide that is strongly anchored to subsite 1 (Figure 2C) and was practically not unbound throughout the whole simulations. This is a notable result that suggests WPAF as being a strong binder of Clec9A, even considering its short size. It is thus suggested that the building blocks of the NPs could be decorated with either the full-length WH peptide, or the WPRF and WPAF truncated variants to target the Clec9A receptor on DC cells.

In the same manner, pBAE (terminal and lateral) were prepared bearing this mutated peptide (M, WPAF sequence). The features of this peptide are particularly interesting, given that replacement of arginine by alanine completely shifts the GRAVY score towards hydrophobic and removes any possible charged residues (Table 2). Indeed, MTer particles display the smallest size, possibly due to higher hydrophobicity (Figure 4-A). It has been well documented that adding hydrophobic moieties to polymer formulation improves the packaging capacity of nucleic acids, thus reducing their size ^51^. MTer also exhibits the lowest zeta-potential (Figure 4B). This is explicable because it is the only peptide with no basic amino-acid residues, while replacing 10% of CH3 end-capped polymer. Regarding encapsulation efficiency, this was slightly lower for MLat (EE 55%, Figure 4-C), while EE was preserved for MTer as compared to the rest of the tested peptides.

Considering previous results, the best improvement could probably be expected in the terminal position. Determination of transfection degrees of the polymer with the new peptide was performed along with an uptake experiment, where CH3-C6 polymer of the particles was labelled with Cy5, and entry of particles and GFP expression in cells was followed by flow cytometry after 2, 6, and 24 hours of incubation. Figure 7A shows that already at 2h time point, most of the particles have entered the cells and reached a plateau (around 90%) and are slowly degrading (80-95% of the polymer is still present after 24h). By hour six, expression of GFP starts to appear, and the formulations that have the best final transfection rate (WHLat and MTer) already reach almost 10%.

**Figure 7.**
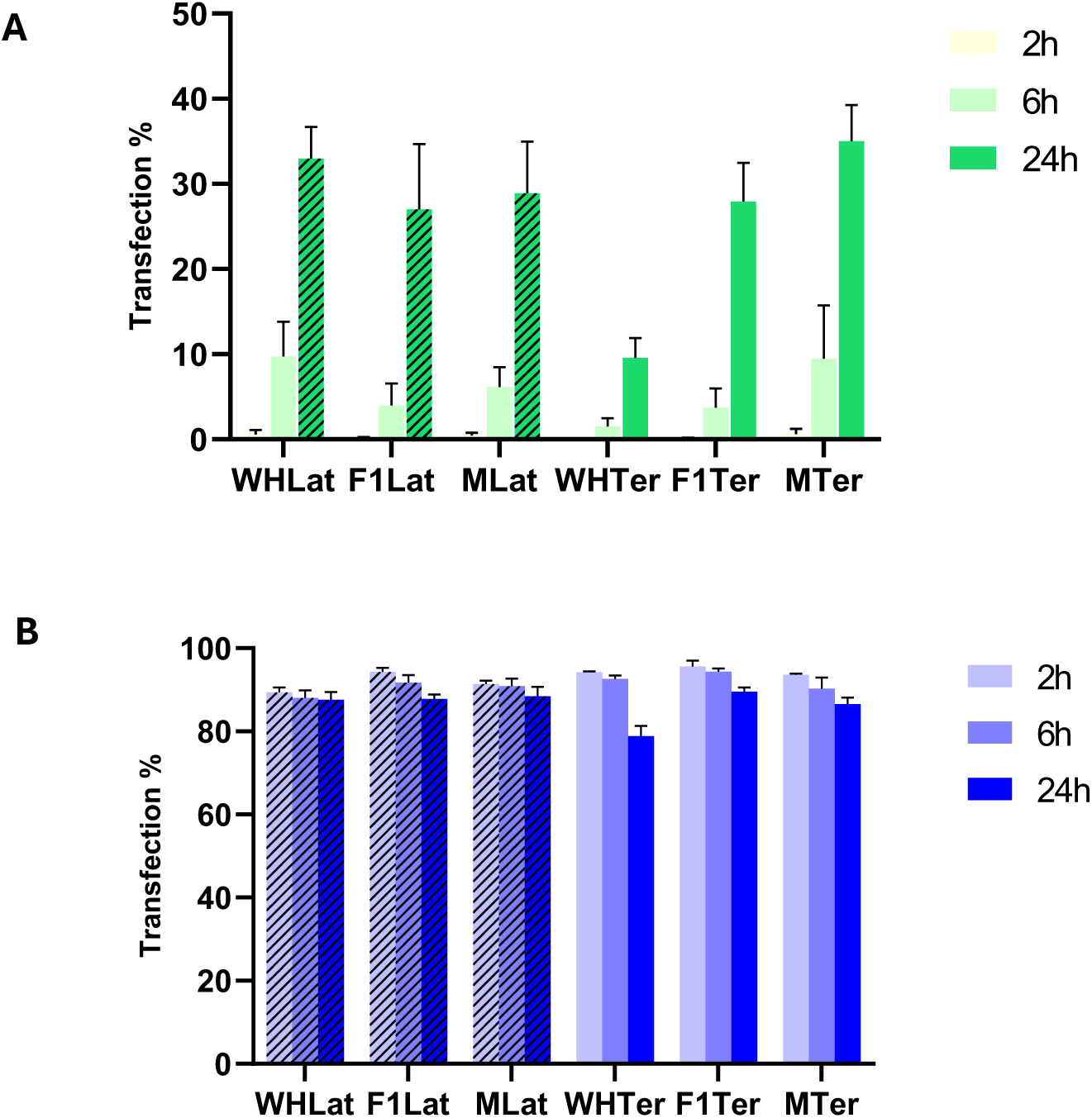
A) GFP expression percentage in THP1 across the targeting formulations at different time points (2,6 and 24h) of incubation B) Percentage of THP1 cells with the presence of Cy5 labelled polymer at distinct incubation times

Analysing the relationship between WPAF and the other two peptides, an improvement in targeting can be observed at all positions, even though in the lateral position, WH still transfects better than both peptides. As previously argued, the expectation that terminal WPAF transfects better than the lateral was proven true, and this is also the best-performing formulation in the experiment.

## Conclusions

In this work, we have presented a unified experimental and computational approach to study the binding of the WH peptide towards the human Clec9A receptor and the dendritic cells’ transfection capacity of NPs grafted with variants of this peptide. The WH peptide, originally designed to target mouse Celc9a, exhibits strong adherence to the human Clec9a counterpart. Based on these studies, two new shortened peptides were designed: WPRF, which retains affinity to the receptor, and WPAF with higher affinity for the receptor. Afterwards, it was confirmed in *in vitro* transfection experiments that all of these peptides improve the targeting capacity of pBAE NPs. The effect of the targeting position of the peptides within a pBAE polymer was found to be dependent on peptide characteristics. This gives a valuable insight when designing future targeting pBAE polymers for biomedical applications. Additionally, this study opens the possibility of using shorter peptides for targeting DC via Clec9a, offering many advantages compared to longer chains, including improved synthesis efficiency (lower costs with higher yield and lower purification burden), chemical stability, and lower steric hindrance when attached to larger structures.

## Supporting information

www.iqs.url.edu

## Acknowledgements

This research was supported by PID2021-125910OB-I00 and PID2024-159704OB-I00 funded by *MCIN/AEI/10.13039/50110 00110 33*, by “ERDF A way of making Europe, by the *Instituto de Salud Carlos III* (ISCIII) (AC22/00042) and by FCAECC (TRNSC213882FORN), both ISCIII and FCAECC from the Joint Transnational Initiative 2021 ERA-NET TRANSCAN-3, European Commission is acknowledged. CF acknowledges the support of *Generalitat de Catalunya*, *Agència de Gestió d’Ajuts Universitaris* (AGAUR) (2021 SGR 00357) and the Departament de Recerca i Universitats de la Generalitat de Catalunya, in the frame of the ICREA Acadèmia 2024. We also thank Andrea Díaz-Tendero for polymer synthesis.

