## Supplementary material for "Targeting dendritic cells with RNA-loaded nanoparticles grafted with short peptides": www.iqs.url.edu

**Supporting information**

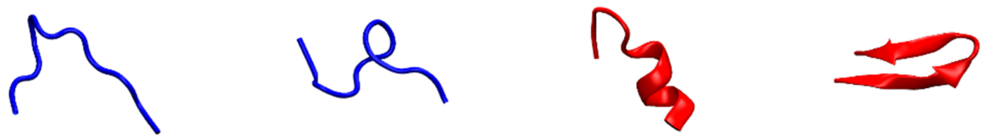

**Figure S1.** Four most representative conformations of WH peptide obtained by structural modelling and molecular dynamics. Blue: structures modelled with PEPFOLD. Red: structures modelled with AlphaFold.

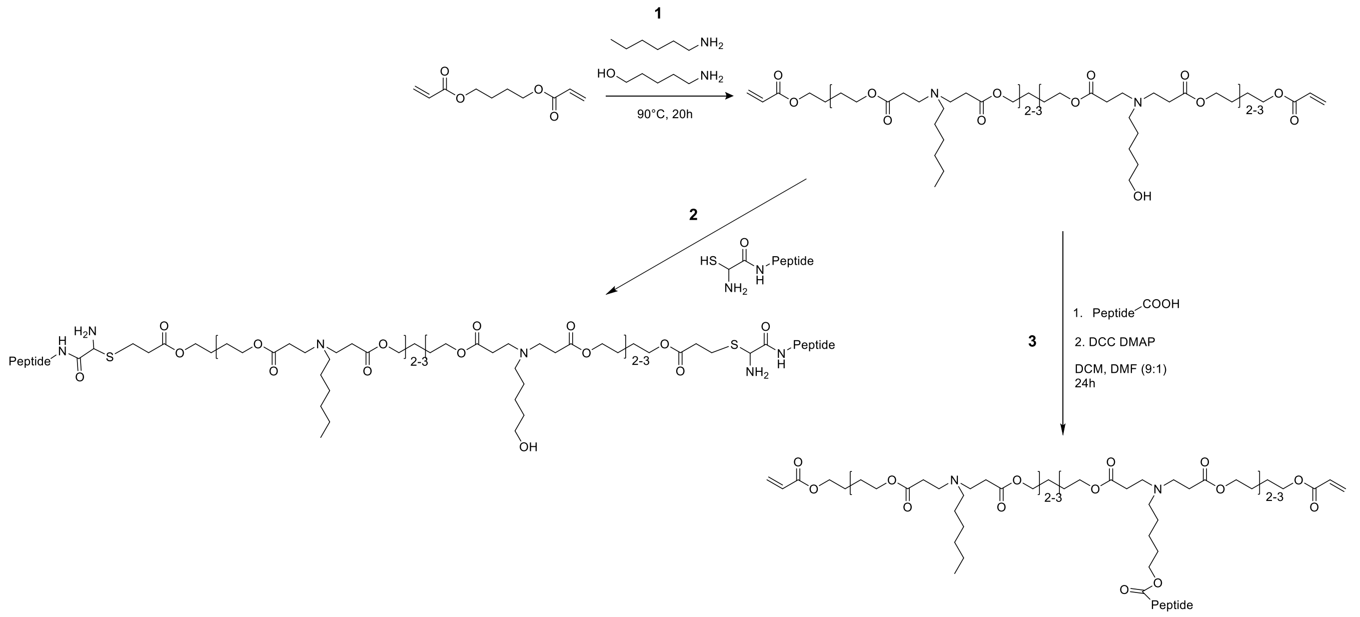


**Figure S2.** Scheme of synthesis reactions of pBAE polymers. **1)** Synthesis of C6 pBAE polymer **2)** Thiol Michael addition to attach cysteine containing peptides to acrylate terminals **3)** Steglich esterification to attach peptides to the lateral chains containing hydroxyl group

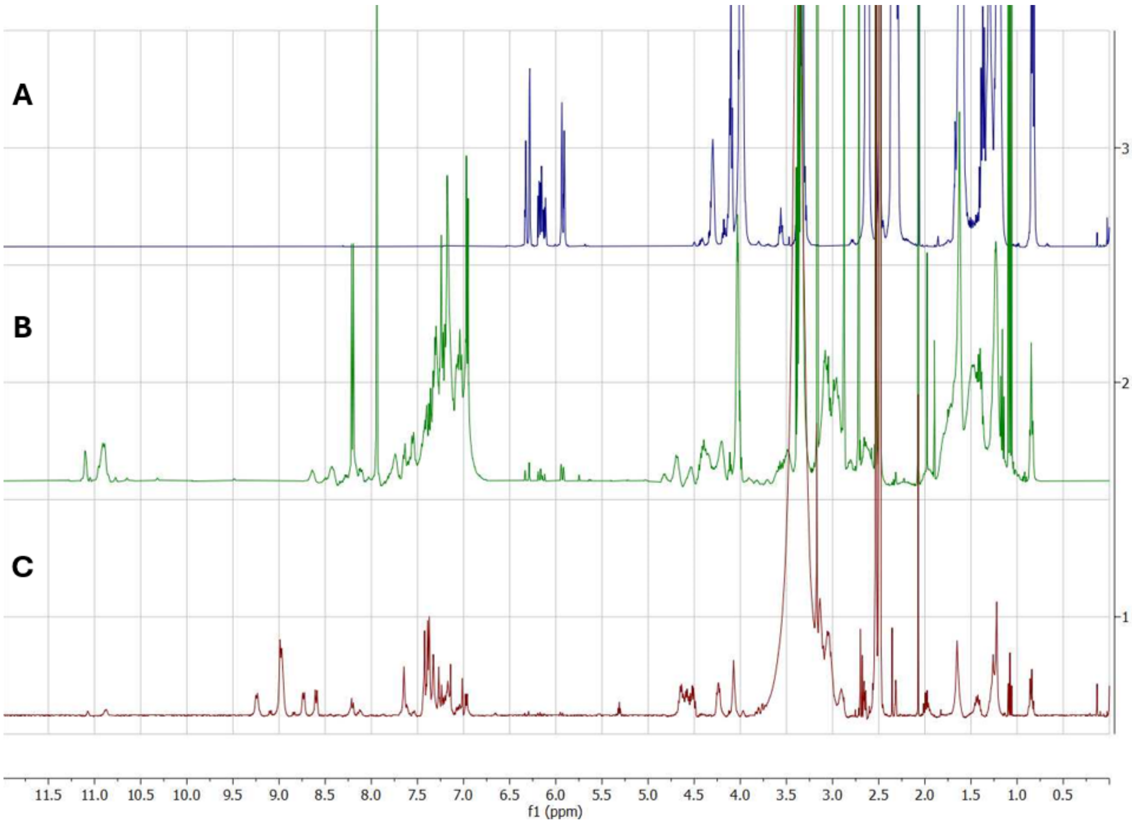


**Figure S3.** Representative NMR spectra of **A)** C6 pBAE **B)** C6 pBAE with laterally attached peptide after Steglich reaction – characterized by apparition of peptide signals **C)** Polymer from (B) then undergoes Thiol-Michael reaction with CH3 peptide – characterized by disappearance of acrylate signals 5.8-6.4 ppm and apparition of histidine aromatic signals

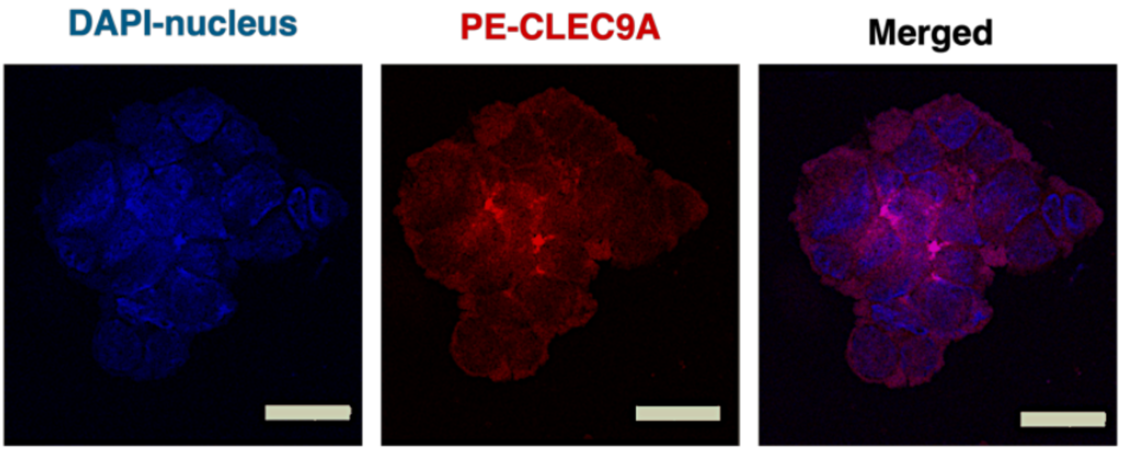


**Figure S4.** CSFM images of THP1 cells. DAPI staining was used to visualize the nucleus and PE-Clec9a antibody to label Clec9a receptor on the surface of the cells. Scale bar = 50 μm.

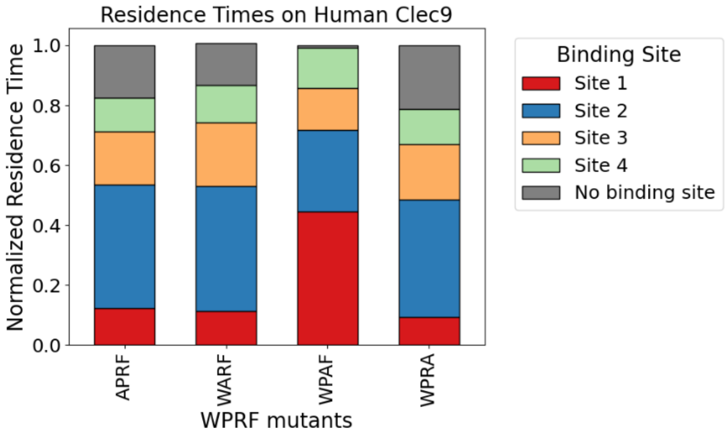


**Figure S5.** Residence time of alanine mutants of the WPRF partial peptide at each binding site.
